# Domains of RNA-binding protein Ssd1 required for tolerating chromosome amplification in yeast

**DOI:** 10.64898/2026.09.25.754501

**Authors:** Adam Jochem, Jamie M. Ahrens, Emma Mayhew, Audrey P. Gasch

## Abstract

Many RNA-binding proteins play pleiotropic roles in the cell, often through disparate domains that can be decoupled. Here we undertook a mutational analysis of yeast RNA-binding protein Ssd1 to better understand domains and processes required for tolerating extra chromosomes, a special class of aneuploidy. Ssd1 is important for cellular fitness when cells carry extra chromosomes, for reasons that are not clear. Ssd1 has multiple distinct domains, many of which are poorly characterized, including multiple RNA binding folds, nuclear import and export signals, an intrinsically disordered domain, an RNA polymerase II-interacting region, and many phosphorylation sites. We measured multiple phenotypes, including the ability to tolerate an extra copy of yeast chromosome XII as a representative aneuploid. Mutating RNA binding domains disrupts fitness in aneuploid, but not euploid, cells and ablates binding of a representative Ssd1 target. However, mutations affecting phase separation or interaction with the kinase Cbk1, which regulates Ssd1 granule formation during the cell cycle, did not affect aneuploidy tolerance. Evaluating correlations across these and other growth phenotypes informed on functional relationships among domains. We discuss implications for Ssd1 function and its role in aneuploidy tolerance in yeast.

## INTRODUCTION

A hallmark of RNA-binding proteins (RBPs) is that many participate in multiple functions in the lifecycle of an RNA. For example, yeast DEAD box helicase Dhh1 plays roles in translation, decapping, and decay, whereas exonuclease Xrn1 contributes to transcription, translation, and co-translational RNA degradation (Blasco-Moreno et al., 2019; Chattopadhyay et al., 2022; Haimovich et al., 2013; Vijjamarri, Gupta, et al., 2023; Vijjamarri, Niu, et al., 2023; Zeidan et al., 2018). How RBPs can function pleiotropically is an interesting topic in studying modular design. In some cases, seemingly disparate processes relate to the same biochemical activity that is utilized in multiple steps of mRNA control (*e.g.* helicase activity that affects many downstream processes). In other cases, activities arise through modular architecture of independently functioning domains (Hentze et al., 2018; Lunde et al., 2007). For example, yeast RBP Pat1 is an mRNA decapping activator that is also involved in transcription, translation, mRNA decay, as well as kinetochore and centromere functions. Several of these Pat1 functions can be at least partly decoupled through domain mutagenesis (He & Jacobson, 2023; Mishra et al., 2013, 2015; Pilkington & Parker, 2008; Pulido et al., 2024a; Wang et al., 1996). Understanding domain architecture can inform on RBP functional evolution and help to isolate the specific domains and processes that drive distinct phenotypes of pleiotropic RBPs.

Ssd1 is an enigmatic RBP that has been linked to many different phenotypes in yeast. One of those phenotypes is sensitivity to chromosome amplification, a specific class of karyotype imbalance or aneuploidy. Euploid cells lacking *SSD1* grow like wild type in the absence of stress, but cells display a major growth defect when individual chromosomes are amplified (Hose et al., 2020; Rojas et al., 2024). Ssd1 was first implicated in aneuploidy tolerance through comparative analysis with aneuploidy-sensitized laboratory strain W303, which carries a hypomorphic truncation of *SSD1* (Hose et al., 2020). Although Ssd1 is necessary to tolerate most duplicated chromosomes, its exact role is not clear. Ssd1 was originally isolated as a second-site suppressor of TORC1 inhibitory phosphatase Sit4 and Protein Kinase A (PKA) regulatory subunit Bcy1 (Sutton et al., 1991). It binds at least 300 mRNAs enriched for those encoding cell-wall proteins, cell-cycle regulators, lipid metabolizing enzymes, and splicing factors, as well as other transcripts involved in diverse processes (Bayne et al., 2022; Hogan et al., 2008; Hose et al., 2020; Jansen et al., 2009). Beyond functional enrichments, mRNAs bound by Ssd1 also have a higher degree of secondary structure, are enriched for mRNAs with upstream uORFs, and encode proteins with a higher degree of disorder (Hose et al., 2020; Dutcher et al., 2024; Spealman et al., 2025). Ssd1 is best (but still relatively poorly) understood for its role in translation and mRNA localization during the cell cycle (Jansen et al., 2009; Kurischko, Kim, et al., 2011; Wanless et al., 2014). Direct phosphorylation of Ssd1 by mitotic regulatory kinase Cbk1 is proposed to alleviate translational repression of cell wall mRNAs required for cellular division. In the absence of Cbk1 kinase activity or when Cbk1-phosphorylated residues in Ssd1 are replaced with alanines, Ssd1 and its targets localize to phase-separated P-bodies (PB), thereby reducing association of those mRNAs with polysomes. Ssd1 has also been implicated in delivery of specific mRNAs to sites of polarized growth (Kurischko, Kim, et al., 2011; Hose et al., 2020). It remains unclear which of these features and processes contribute to aneuploidy tolerance.

Ssd1 is a large protein of 1250 amino acids spanning several domains, including an RNase-II-type binding domain and multiple OB folds that recognize single-stranded nucleic acids. It is orthologous to mammalian exonuclease Dis3L2 that degrades poly-uridylated RNAs (Luan et al., 2019); however, Ssd1 lost the ability to bind cations required for catalysis and shows no nuclease activity (Bayne et al., 2022; Uesono et al., 1997). Its amino terminus is largely unstructured and includes a glutamine (Q) and asparagine (N)-rich intrinsically disordered region (IDR) peppered with phosphorylation sites, including those recognized by Cbk1, Cdc28, and PKA as well as other kinases (Gógl et al., 2015; Myers et al., 2019; Oughtred et al., 2021). IDRs of other RBPs contribute to phase separation, which can be modulated by phosphorylation (J. Li et al., 2022; M. Li et al., 2023; Ramachandran & Potoyan, 2025). The unstructured amino terminus also contains a domain shown to interact with phosphorylated carboxyl-terminal domain (CTD) of Pol II. The consequence of that interaction is not known, but Ssd1 has been identified as a suppressor of polymerase mutations (Chiannilkulchai et al., 1992; Stettler et al., 1993). Consistent with nuclear and cytoplasmic functions, Ssd1 has predicted nuclear-localization (NLS) and -export (NES) sequences (Kurischko, Kuravi, et al., 2011; Kurischko & Broach, 2017). How these domains interface with one another is not known.

To better understand the domains of Ssd1 important for aneuploidy tolerance, we generated a panel of 20 mutants interrogating domains, residues, and motifs, and then scored mutant phenotypes across multiple environments and functions. We studied each mutant in euploid cells and aneuploid cells with an extra copy of Chromosome XII (Chr12) in wild oak-soil isolate YPS1009. Chr12 duplication is among the chromosome amplifications most dependent on Ssd1 for fitness. Ssd1 dependence is well modeled by protein-coding genes on the amplified chromosome, including those that are highly toxic when duplicated individually in the *ssd1Δ* euploid (Dutcher et al., 2024; Rojas et al., 2024). However, deciphering direct from indirect effects in aneuploid cells has been a challenge. Here we interrogated domains involved in RNA binding, granule formation, Cbk1 interaction, tolerance of cell-surface stress and rapamycin, and ultimately tolerance of Chr12 duplication. Our results suggest that RNA binding is essential for Chr12 tolerance, while granule formation and domains for Pol II and Cbk1 interactions are dispensable. Several functional Ssd1 variants displayed unexpected genetic interactions with *CBK1* deletion. Together, these results inform on mechanisms by which Ssd1 endows cells with aneuploidy tolerance.

## RESULTS

We annotated the Ssd1 sequence based on previous reports in the literature (Jansen et al., 2009; Kurischko, Kim, et al., 2011), comparison to orthologous protein structures available at the time, and conservation across fungal lineages (see Methods, Dataset 1). A summary of these domains is illustrated in **Fig 1**. We designed mutations to test these domains, introducing large deletions of the amino terminus (mutants C and H, **Fig 1**) or carboxyl terminus (A, B, O, and Q), small scale deletions or substitutions that remove conserved domains (I and M), and substitutions predicted to disrupt nuclear import/export (K, L, T) or RNA binding, based on the crystal structure of the mouse Dis3L2 ortholog available at the time (I, J, N, P, R, S) (Powell et al., 2025). We also tested the impact of the amino-terminal IDR, by wholesale deletion (C, H) or by substituting asparagines with leucines (NtoL, mutant D) and or by changing glutamines to alanines (QtoA, mutant E). To interrogate Cbk1 functions, we ablated Cbk1 docking sites (mutant G) and generated a phospho-mimic of 4 Cbk1-phosphorylated residues (at sites 40, 42, 160, 164, mutant F). Each mutant was fused to an in-frame GFP tag with native *SSD1* 3’ UTR and introduced into the genomic *SSD1* locus in euploid or YPS1009 with an extra copy of Chr XII (YPS1009_Chr12).

**Figure 1.**
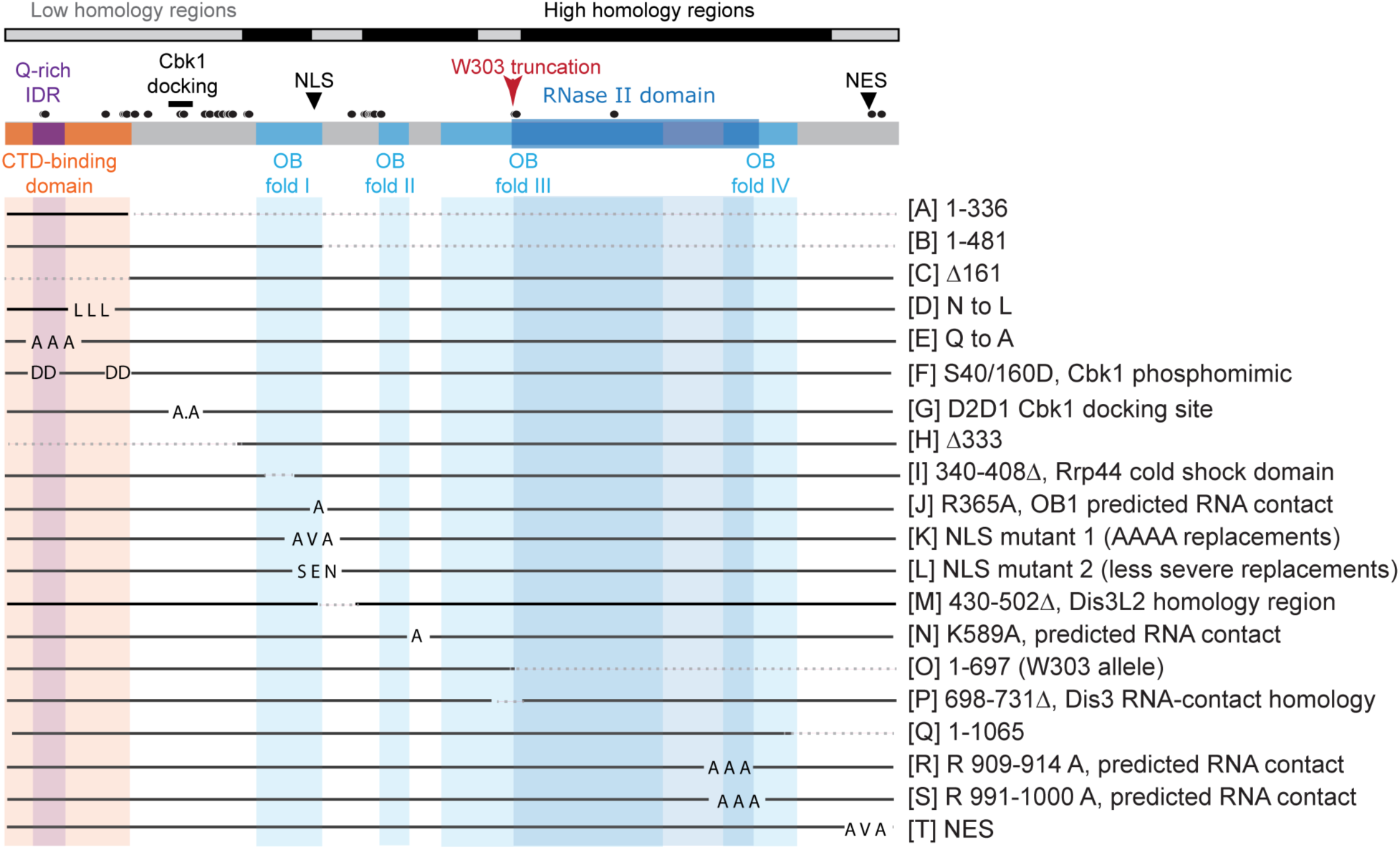
Domains and mutants of Ssd1. The top panels annotate domains and regions of Ssd1, see text for details. Circles, phosphorylation sites in Biogrid (Oughtred et al., 2021); CTD, Pol II carboxyl-terminal domain; Cbk1, Cbk1 docking sites; W303 truncation, location of premature stop codon in the W303 strain; dashed lines, deletions. The diagram below represents the position and type of substitutions or deletions, which are annotated with mutant name (A-T) and short description to the right. Detailed mutant descriptions and domain annotation are available in Table 1 and Dataset 1.

### Mutant phenotyping implicates domains important for Chr12 tolerance

We began by measuring the ability of each mutation to support Chr12 duplication. In rich medium, YPS1009_Chr12 grows at ∼85% the rate of the euploid wild type, whereas the growth rate of *ssd1Δ* YPS1009_Chr12 is reduced to ∼60% of wild-type or *ssd1Δ* euploids (Hose et al., 2020). None of the mutants grew significantly differently from wild type when expressed in the euploid, as expected since the euploid *ssd1Δ* has no growth defect in this background (p > 0.05, **Fig S1**). However, the mutants produced significantly different growth rates when expressed in the YPS1009_Chr12 aneuploid (**Fig 2**). We classified aneuploid mutants that grew indistinguishably from wild-type or from *ssd1Δ* aneuploids (black or grey bars, respectively, **Fig 2**) versus those with moderate (dark blue) or more severe (light blue) partial growth defects (p<0.05, replicate-paired T-test). We verified Ssd1 protein abundance of some of the most impactful mutants, and observed that several deletion mutants (M and P) and NLS mutant L that behaved like the *ssd1Δ* aneuploid produced little soluble protein, suggesting stability / solubility defects (Kurischko et al., 2017) (**Fig S2**).

**Fig 2.**
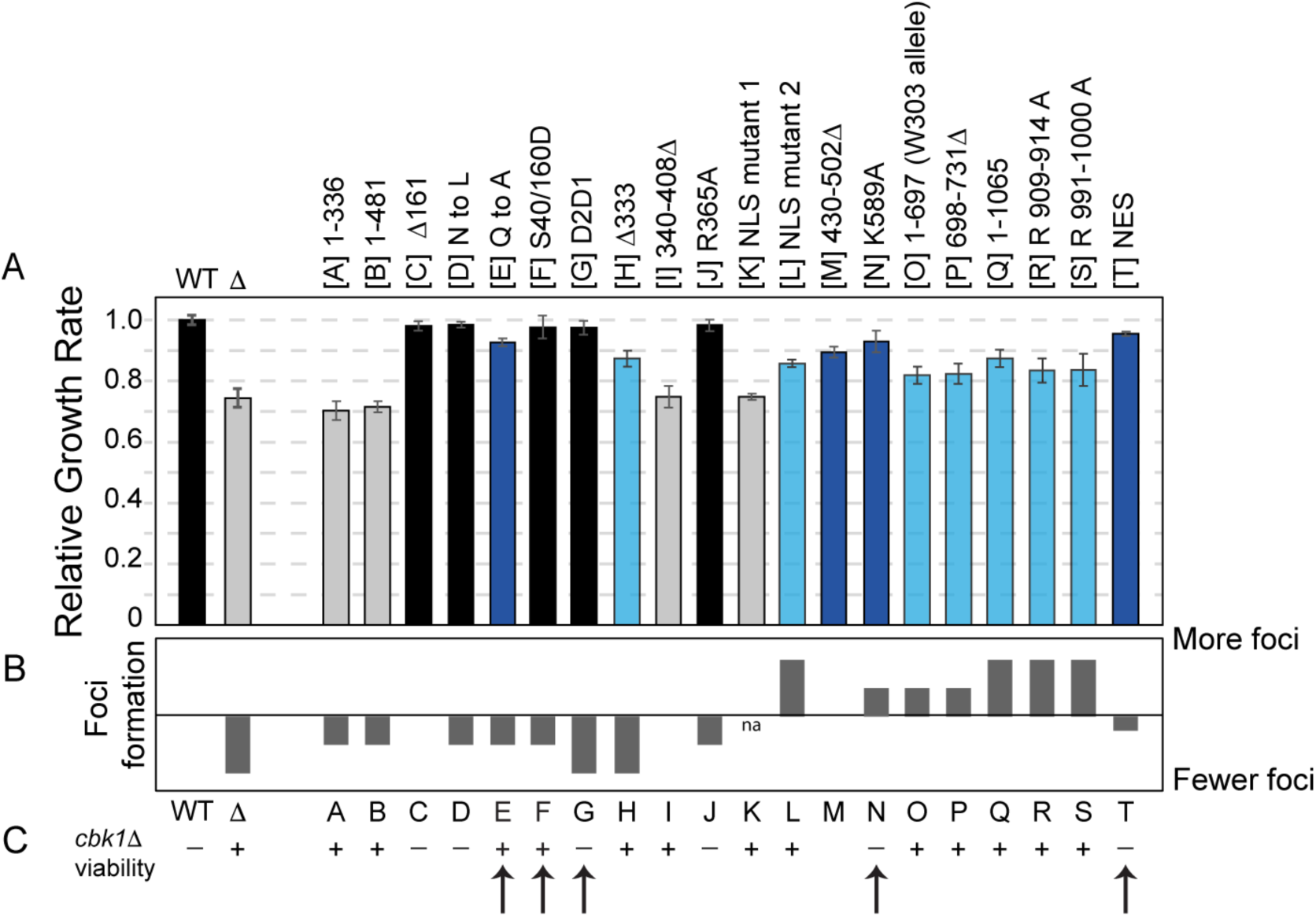
Tolerance of Chr12 duplication in ssd1 mutants. A) Average and standard deviation (n >= 3) of growth rates of each indicated strain, relative to the wild-type strain. Histograms are colored by growth defect: black, statistically similar to wild-type (p > 0.05, replicate-paired T-test); grey, statistically similar to *ssd1Δ* (Δ) aneuploid (p>0.05); light blue and dark blue indicate statistically significant defects compared to wild-type (p < 0.05) that were more (<90% wild-type growth rate) or less (>90% growth rate) severe. B) Semi-quantitative analysis of % cells with foci upon glucose starvation, see Methods. C) Binary assessment of mutant viability in euploid *cbk1Δ* cells (see Fig S4). Arrows indicate cases discussed in the text.

As expected, the gross deletions in mutants A and B, ablating all but the first 336 or 481 residues, produced a null phenotype. Interestingly, these mutants were expressed at 5-15X higher levels in both euploids and aneuploids. It is possible that sequences governing the suppression of Ssd1 abundance have been deleted, a hypothesis what will require further experimentation. Other mutants were more informative. Most mutations in the amino-terminal quarter of the protein had little effect on aneuploid growth rate, including deletion of the Pol II CTD-interacting domain (mutant C), substitution of asparagines in the IDR (D), replacement of Cbk1 docking sites (G), or substitutions mimicking Cbk1 phosphorylation at positions 40, 42, 160, 164 (F, **Fig 2**, black bars). An interesting exception was the QtoA mutant: replacing glutamines in the IDR produced a moderate defect in Chr12 tolerance (**Fig 2**, dark blue bar), even though wholesale deletion of the region produced a wild-type phenotype (**Fig 2**, mutant E versus C). On closer inspection, we realized that mutant C is expressed ∼8-fold higher in the Chr12 aneuploid compared to the euploid (p = 0.047, paired T-test, **Fig S2**), raising the possibility that over-expression complements a functional defect (see more below). Deletion of the amino-terminal 333 residues (mutant H), which abuts the first OB1 fold, produced a more pronounced growth defect than did the other N-terminal mutants (**Fig 2**).

In contrast to the amino-terminal portion, which was generally not required for aneuploidy tolerance, alteration of the carboxyl-terminal half of the protein, including RNA binding domains, induced more pronounced phenotypes. All but one of the mutants predicted to affect RNA binding (including mutants N, P, R, S in OB domains and deletion mutants ablating the RNase II domain) produced a severe, but not quite null, aneuploid growth defect (**Fig 2**, light blue bars). The exception was R365A in domain OB1 (mutant J), which produced no growth defect. This mutant was expressed substantially higher in aneuploid versus euploid cells (p = 0.022, paired T-test, supplementary **Fig S2**), raising the possibility of complementation. Together, these results suggested that RNA binding is important for Chr12 tolerance. Unfortunately, we were unable to interrogate the role of nuclear localization, since neither NLS-2 nor NES mutants showed clear differences in localization by z-stack microscopy.

### Binding of SUN4 mRNA correlates with Chr12 tolerance

To test the hypothesis that RNA binding is important for aneuploidy tolerance, we quantified RNA binding capability of each mutant expressed in euploid cells using RNA immunoprecipitation. We chose euploid cells to study inherent binding capabilities, separate from altered protein abundance observed in several mutant aneuploids. We characterized enrichment of representative Ssd1 target *SUN4* versus non-target *ACT1* (**Fig 3**). Most of the mutants predicted to disrupt RNA binding recovered significantly less *SUN4*. The exception was mutant R365A in OB fold 1 (mutant J), which appears to bind wild-type amounts of *SUN4*. We conclude that substituting arginine at 365 does not disrupt Chr12 tolerance or RNA binding, at least of *SUN4*. Despite a noisy assay, all mutants with a growth defect in Fig 2 showed a defect binding *SUN4*. This supports the model that RNA binding is important for Ssd1’s role tolerating extra chromosomes. Yet most of these mutants had some tolerance of Chr12 duplication, greater than that of the *ssd1Δ* strain. It is possible that these mutants retain some RNA binding activity that is not apparent in this experiment (including non-specific RNA binding reported for Ssd1 (Uesono et al., 1997)), or that Ssd1 activities beyond RNA binding contribute weakly to tolerance of Chr12 duplication.

**Figure 3.**
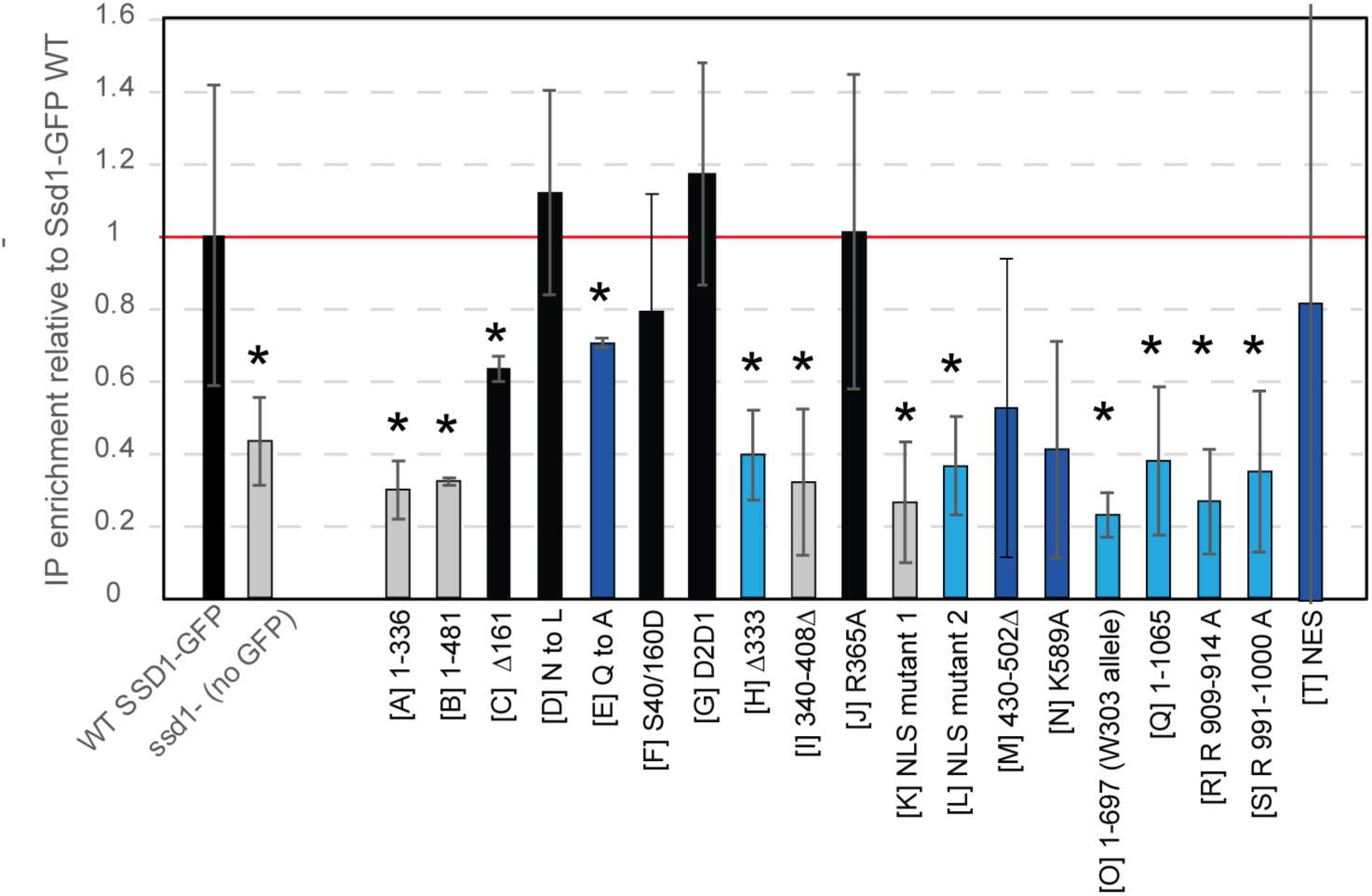
Ssd1 binding of *SUN4* mRNA correlates with Chr12 tolerance. The average and standard deviation (n = 2-3 for mutants, >7 for controls) of immunoprecipitation (IP) enrichment over input control was assessed (see Methods). Asterisk, p<0.05, one-tailed paired T-test versus wild-type control. Bars are colored by the growth category of each strain as described in Fig 1 for comparison.

Somewhat surprisingly, QtoA mutant E that has a defect tolerating Chr12 aneuploidy showed reduced *SUN4* recovery in the IPs, consistent with weaker binding. This was surprising, since these substitutions are far in primary sequence from the predicted RNA binding domains (**Fig 1**) and instead fall in an Alpha-fold predicted helix within the unstructured region of Ssd1 (Bayne et al., 2022; Jumper et al., 2021). Mutant C lacking the first 161 residues, including RNA Pol II-interacting domain, also had a partial defect binding *SUN4* when expressed in euploid cells, similar to the QtoA mutant (**Fig 3**) – but it had no defect tolerating Chr12 duplication (**Fig 2**). One possibility is that the increased abundance of mutant C in aneuploid cells complements the partial binding defect of the protein. Nonetheless, these results show a high correlation between the ability to bind *SUN4* RNA and the ability to tolerate Chr12 duplication.

### Granule formation is not required for Chr12 tolerance

Ssd1 can phase-separate into granules, including PB induced by stress or *CBK1* inhibition (Kurischko, Kim, et al., 2011; Kurischko & Broach, 2017; Xing et al., 2020). Past work from our lab identified other RBPs that contribute to Chr12 aneuploidy tolerance, albeit much less-so than Ssd1 (Dutcher & Gasch, 2024). Because many of these associated factors localize to PBs, it is plausible that the phase separation of Ssd1 plays a role in mediating aneuploidy tolerance. We therefore measured the propensity of ssd1 mutants to form GFP foci upon glucose starvation to induce PBs. Ssd1 can be readily identified in PB when over-expressed or in cells predisposed to PB formation (Kurischko, Kim, et al., 2011; Xing et al., 2020). In our hands, few wild-type cells (10-20%) harbored Ssd1-GFP foci, typically one per cell, when starved for glucose (see Dataset 3); nonetheless, it was enough to compare mutants semi-quantitatively (see Methods).

Although most mutations in the IDR region did not affect Chr12 tolerance, many of those mutants produced fewer foci when aneuploid cells were starved for glucose (**Fig 2B**). The largest deficit was seen when Cbk1 docking sites were ablated (mutant G), producing almost no cells with foci (**Dataset S3**). This result was unexpected, since the loss of Cbk1 interactions was hypothesized to localize Ssd1 to granules (see more below). Loss of the entire N-terminal 333 amino acids (mutant H) also produced a major deficit in foci production, while other mutations in this region produced more subtle deficits. The only exception was mutant C that produced a similar number of foci to wild type, perhaps due to increased ssd1 protein abundance (**Fig S2**). Together, these results are consistent with the model the IDR is important for phase-separation. However, there was no correlation between the ability to form foci and Chr12 aneuploid growth rate (**Fig 2A**). This was most clear for the Cbk1 docking mutant that produced no foci and showed no sensitivity to Chr12 duplication. We conclude that the ability to form visible foci as seen during carbon starvation is not required to tolerate Chr12 duplication. We note that it is possible that phase separation in response to aneuploidy, too subtle to view by microscopy, could be governed by different domains.

To further probe the role of Cbk1 interactions, we tested the genetic interaction of these mutants with Cbk1, by measuring their viability when expressed in a euploid *cbk1Δ* strain (**Fig 2C**). Wild-type *SSD1* is lethal when expressed in *cbk1Δ* cells, whereas *ssd1Δ cbk1Δ* cells are viable (Jorgensen et al., 2002; Kurischko et al., 2005). For the most part, viability of these mutants tracked Ssd1 function as assessed by Chr12 tolerance: *ssd1* mutants tolerant of Chr12 duplication were lethal when expressed in *cbk1Δ* cells, whereas *ssd1* mutants with a growth defect in the Chr12 aneuploid were viable in a *cbk1Δ* background. Two of the mutants with intermediate Chr12 sensitivity (N and T, Fig 2 arrows) were not viable in *cbk1Δ* cells, suggesting they maintain enough functionality to be lethal without Cbk1 inhibition. However, there were a few exceptions. Mutant F that mimics the Cbk1-phosphorylated state showed full tolerance of aneuploidy and viability when expressed in *cbk1Δ* cells, as expected if Cbk1 is no longer required to phosphorylate those sites. Mutant G lacking Cbk1 docking sites was unexpectedly viable in wild-type cells. This was surprising, since the introduced mutation was previously shown to ablate Cbk1 binding, which would mimic the *cbk1Δ* state with regard to Ssd1 regulation (Gógl et al., 2015). Mutant G showed no sensitivity to Chr12 duplication, and thus like the wild-type strain was not viable in the *cbk1Δ* background. It is possible that Cbk1 can still bind the mutant in this strain background, and that effects of mutant G regulate phase separation by other means beyond Cbk1 control. QtoA mutant E was also unexpectedly viable without Cbk1, whereas other mutants with comparable Chr12 tolerance were lethal. These results hint that aspects of Cbk1 regulation of Ssd1 remain to be elucidated (see Discussion).

Many RBPs require RNA binding to phase separate, perhaps due to generalizable biophysical requirements (Fuller et al., 2020; Roden & Gladfelter, 2021; Van Treeck et al., 2018; Zhang et al., 2015). Thus, we predicted that mutants with a *SUN4* binding defect may produce fewer foci. We instead observed that these mutants produced more foci, often significantly larger in size (**Fig 2B** and **S3**). Loss of RNA binding causes some RBPs like FUS and TDP43 to form solid cytoplasmic structures, rather than fluid phase transitions, suggesting that RNA binding solubilizes these RBPs in the cell (Maharana et al., 2018). It is possible that Ssd1 falls into this class and that RNA binding is required to maintain granules in a fluid state, although other models could also explain the effect.

### Correlations across mutant phenotypes point to functional requirements for Chr12 tolerance

To further understand the determinants of Chr12 tolerance, we measured several other phenotypes associated with Ssd1 function or aneuploidy tolerance. These included growth under cell wall stress induced by calcofluor white, tolerance of membrane stress induced by SDS detergent, the ability to respire on glycerol (since YPS1009_Chr12 *ssd1Δ* is sensitive to non-fermentable carbon sources (Hose et al., 2020)), and sensitivity to TORC1 inhibitor rapamycin. We measured phenotypes in euploid and aneuploid mutants and then compared phenotype distributions across the set of mutants (**Fig 4A**), generating a heatmap of pairwise correlations (**Fig 4B**). Sensitivity to Chr12 duplication (measured as aneuploid growth rate on glucose) was most correlated with sensitivity to SDS, whether measured in euploid or aneuploid cells. It was also correlated with aneuploid growth defects on glycerol. The ability to bind RNA and sensitivity to cell wall stress or rapamycin were also highly, but slightly less well, correlated. These results indicate that domains important for Chr12 tolerance are also important to survive cell surface stress, to bind RNA, and to respire well in the aneuploid state. As predicted from results above, the ability to form granules was not correlated with aneuploid growth rates and was weakly negatively correlated with viability in *cbk1Δ* cells. Analyzing the phenotypic connections suggests relationships across domains and with regard to aneuploidy tolerance, discussed in detail below.

**Figure 4.**
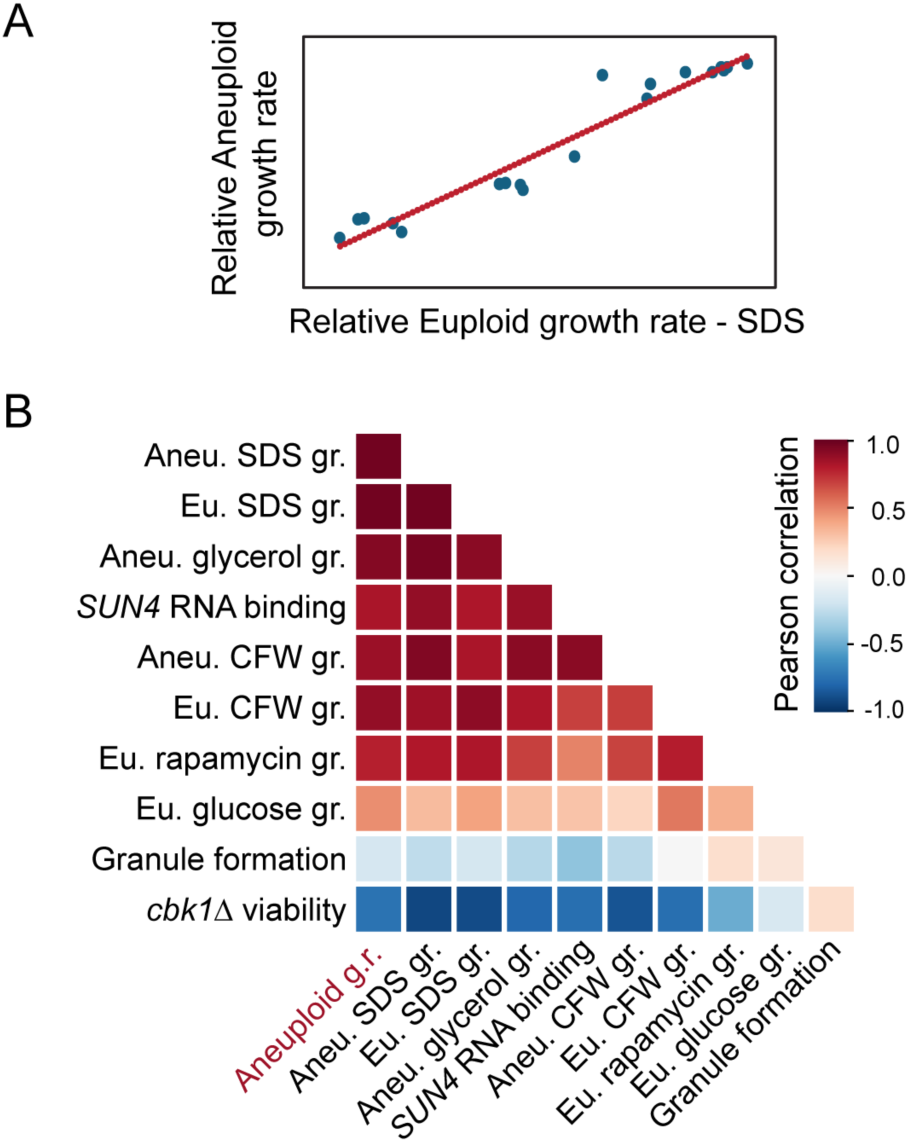
Correlations across mutant phenotypes implicate functions important for Chr12 tolerance. A) Example comparison of mutant phenotypes across two conditions. B) Heatmap of pairwise Pearson correlations for each pair of phenotypes listed, colored according to the key. Growth rate, g.r.

## DISCUSSION

Ssd1 is required for tolerance of most large chromosome duplications in wild yeast strains (Hose et al., 2020; Rojas et al., 2024), but how it enables aneuploid fitness has remained unclear. Past modeling work strongly suggests that the mechanism relates to protein coding genes on the duplicated chromosome (Dutcher et al., 2024; Dutcher & Gasch, 2024; Rojas et al., 2024). Yet there is no evidence that Ssd1 plays a generalizable role in silencing mRNA or protein expression from the duplicated chromosome, and no evidence of a common signature shared amongst all of its bound mRNAs or their encoded proteins (Dutcher et al., 2024; Hose et al., 2020). Part of the challenge in deciphering Ssd1’s role is that *ssd1Δ* aneuploids have widespread secondary responses to their extreme stress, masking direct effects from indirect consequences. Here we took a different approach to studying the role of Ssd1 through domain analysis. The results suggest new information about this RBP while refining models for its role in aneuploidy tolerance.

One question at the outset of this project was whether functions of Ssd1 could be decoupled by mutating different domains. Mapping phenotypes onto the primary and secondary protein structures implicates regions important for various Ssd1 properties (**Fig 5**). Most domains predicted to bind RNA disrupted *SUN4* binding when mutated, suggesting that both OB folds and noncatalytic RNAase II domains contribute (**Fig 5A**). The amino-terminal unstructured region containing an IDR is important for visible granule formation, as predicted. Among the most informative mutants was the QtoA substitution within this region, predicted to fall into a helical region of the unstructured IDR (**Fig 5B**). Glutamine-rich regions are often found in nucleic acid binding proteins (Karlin & Burge, 1996; Schaefer et al., 2012). They can mediate protein-protein interactions but also make contacts with RNA (Martinez-Yamout et al., 2023; Moldovean-Cioroianu, 2024; Tan & Frankel, 1998). Substituting glutamine residues reduced granule formation as well as RNA binding, as did deletion of the entire region in mutant C. These results indicate that residues outside of well-known RNA binding domains contribute to RNA binding, as suggested previously (Bayne et al., 2022). Both mutants are predicted to abolish the Pol II CTD-interacting region, whose function remains unknown. An intriguing possibility is that Ssd1 associates with nascent RNA in the nucleus, perhaps via CTD interactions, similar to several other proteins with which Ssd1 interacts (Blasco-Moreno et al., 2019; Chattopadhyay et al., 2022; Kurischko et al., 2017; Pulido et al., 2024b). If so, the interaction is not essential for Chr12 tolerance, since the Δ161 mutant C tolerates Chr12 duplication, perhaps enabled by its increased expression specifically in aneuploid cells.

**Figure 5.**
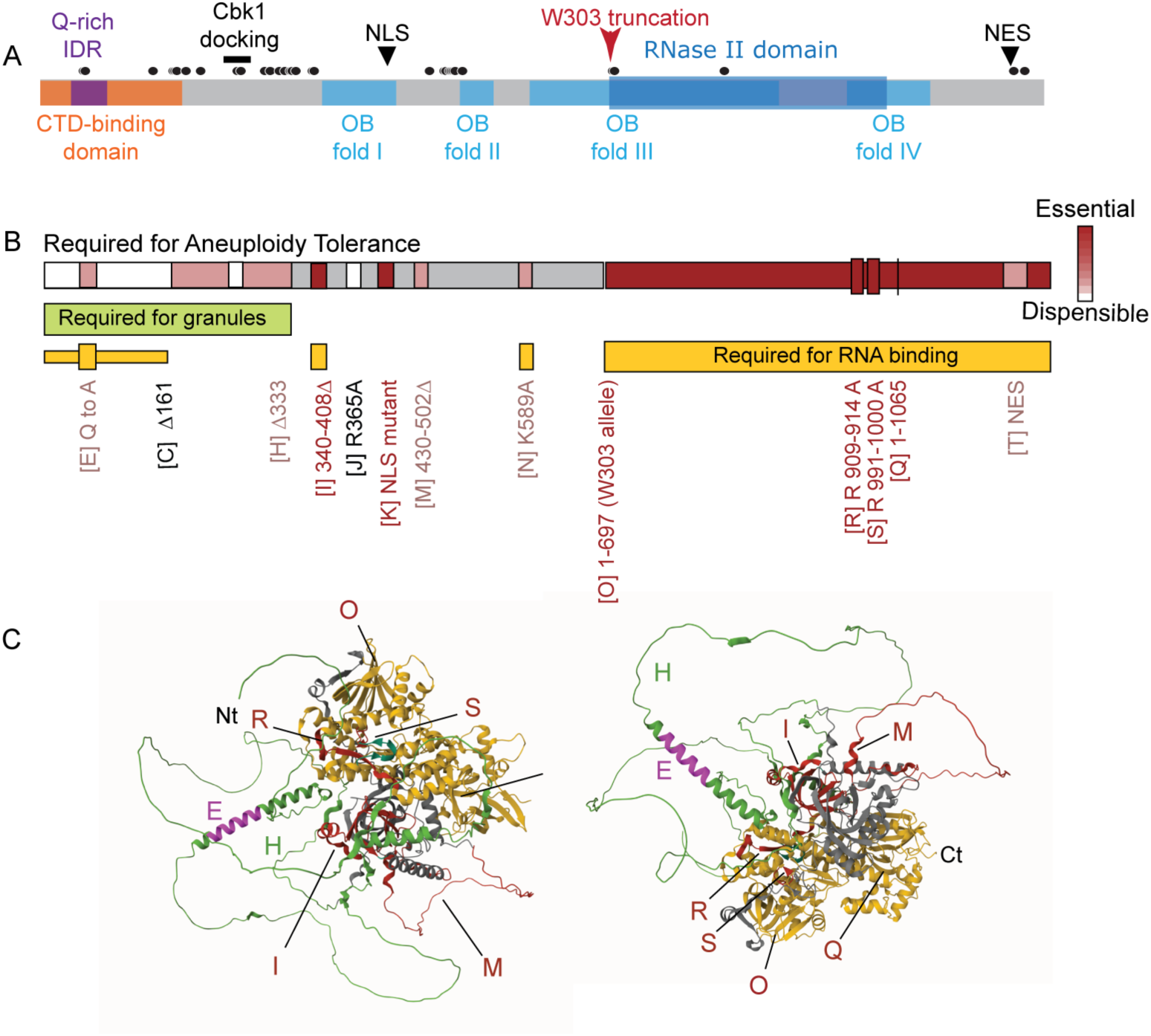
Summary of domain requirements for tolerance of Chr12 duplication. A) Representation of Ssd1 domains from Fig 1 for comparison. B) Domain requirement for Chr12 tolerance, based on informative mutations listed below the figure. Domains required for granule formation or RNA binding are indicated by green or orange boxes (grey, not tested). Mutants that produced little soluble protein were not scored for granule formation or RNA binding. C) Location of informative substitutions or deletions mapped onto the Alpha-fold Ssd1 structure, shown in two orientations. Colors refer to: green, regions important for granule formation; magenta, location of Q residues in mutant E; orange, domains from B important for RNA binding; red, impactful mutants that affect protein folding and/or RNA binding, as indicated by the labeling. The amino-terminal region is unstructured and predicted by Alpha-fold (Jumper et al., 2021). The core of the structure is nearly superimposable with the published Ssd1 structure by (Bayne et al., 2022).

### Domains important for Chr12 tolerance

For the most part, the relative defects of the mutants were well correlated across phenotypes, from tolerance of cell-surface stress and rapamycin treatment to respiratory defects specific to YPS1009_Chr12 (**Fig 4B**). This suggests that Ssd1 properties important for one phenotype are most often required for another. Mutants defective in RNA binding had major defects in most phenotypes, including the ability to form normal granules. These results underscore the importance of RNA binding. Yet, most mutants with gross mutation of RNA binding domains, and consequential loss of *SUN4* mRNA binding, maintained residual tolerance of Chr12 duplication. Ssd1 can bind RNA through sequence-specific interactions and in a sequence-independent manner (Bayne et al., 2022; Uesono et al., 1997). One possibility is that residual RNA binding, perhaps even through protein-protein interactions, supports enough activity for partial aneuploidy tolerance. Another possibility is that Ssd1 supports additional functions outside of RNA interactions, which will require future study to identify.

While RNA binding is important for Chr12 tolerance, the ability to form visible granules is not. This was surprising, both because PB localization is central to the Cbk1 regulation model (Jansen et al., 2009; Kurischko, Kim, et al., 2011) and because past work from our lab implicated multiple PB RBPs in Chr12 tolerance (Dutcher & Gasch, 2024). It is possible that mutants studied here can still phase separate, but fail to congeal into larger granules visible by microscopy (Escalante & Gasch, 2021; Glauninger et al., 2022). But another possibility is that the ability to form granules is dispensable, whereas RBPs that happen to localize to PB are important. Many of the PB factors implicated in aneuploidy tolerance play diverse roles in translation and mRNA decay (Dutcher & Gasch, 2024). Ssd1 and several of these RPBs bind overlapping sets of targets, suggesting they may function together on the same transcripts. Several of those RBPs function in translational regulation, mRNA decay, and translational control during mRNA localization.

Ablating the Cbk1 docking sites was unexpectedly viable, did not affect Chr12 tolerance, and eliminated formation of visible granules during carbon starvation. While mutation of the Cbk1 docking sites was previously shown to disrupt interaction with Cbk1 (Gógl et al., 2015), it is possible that Cbk1 still phosphorylates this mutant. Furthermore, this sequence could have additional roles outside of Cbk1 interaction that contribute to phase separation. Cbk1 functions at the bud neck and in daughter cells, raising the possibility that it regulates only a localized pool of Ssd1 molecules (Colman-Lerner et al., 2001; Mazanka et al., 2008; Weiss et al., 2002).

### Insights into the role of Ssd1 in aneuploidy tolerance

Several possibilities remain for an RNA-dependent role for Ssd1 in tolerating extra chromosomes. Numerous lines of evidence implicate translational control. First, Ssd1 interacts physically with the ribosome and many proteins involved in translation initiation (Engel et al., 2025). Its deletion perturbs translation during cell-cycle progression and during healthy aging (Hu et al., 2018; Jansen et al., 2009; Ohyama et al., 2010; Wanless et al., 2014). Second, both wild-type and especially *ssd1Δ* aneuploids are sensitive to ribosome-stalling drugs, and *ssd1Δ* aneuploid fitness can be augmented by deletion or over-expression of other translation factors or tRNAs (Dutcher et al., 2024; Dutcher & Gasch, 2024; Escalante et al., 2025; Hose et al., 2020; Rojas et al., 2024; Torres et al., 2007). Third, Ssd1 has been linked to translational suppression of single-gene duplicates, including those with upstream uORFs. Indeed, Ssd1-bound mRNAs are enriched for harboring regulatory uORFs (Dutcher & Gasch, 2024; Spealman et al., 2025). Finally, loss of *SSD1* in YPS1009_Chr12 cells increases abundance of several proteins encoded by Ssd1-bound mRNAs, consistent with altered translational control – but only in aneuploid cells, and only at a small subset of Ssd1-bound mRNAs (Hose et al., 2025). If Ssd1 does regulate mRNA translation, it is likely to do so at only some of its bound mRNAs.

Ssd1 could also influence mRNA decay. Some bound mRNAs show increased abundance in aneuploids lacking *SSD1* (Hose et al., 2025). Ssd1 interacts both genetically and physically with multiple mRNA decay factors, including exonuclease Xrn1, decapping factors Lsm4, Dcp2, Pat1, and others involved in mRNA metabolism (Engel et al., 2025). There is no consistent pattern of abundance change across all bound mRNAs when *SSD1* is deleted, suggesting target-specific and perhaps context-dependent effects. While modeling implicates the importance of protein-coding genes, it is notable that Chr12 studied here also carries the rDNA locus, whose role in Chr12 sensitivity is not clear. There are no known links between Ssd1 and RNA processing, but it remains possible that an additional function at rRNA contributes. Although questions about Ssd1 and extra-chromosome tolerance remain, our work provides new information on the functions of Ssd1 domains, peculiarities of phase separation and RNA binding, and the ways in which RBPs can play diverse roles in physiology.

## METHODS

### Strains and growth conditions

Strains used in this study, mutation descriptions, and rationale are listed in Table 1. Unless otherwise noted, YPS1009 euploid or YPS1009_Chr12 aneuploid strains were grown in rich YPD medium to log phase before experimentation. OD_600_ measurements taken over time were fit with exponential curves to attain growth rates. Cells were exposed to drugs added at the start of each growth curve, including: 25 µM calcofluor white (Sigma), 0.008% SDS (Thermo), 0.001 µ/mL rapamycin (Sigma), or shifted to YP 2% glycerol for nonfermentable growth. Growth rate data are available in Dataset 2.

### Sequence analysis and mutant construction

Regions of low and high homology were assessed using COBALT (Papadopoulos & Agarwala, 2007) applied to an alignment of 23 fungal Ssd1 sequences from the Broad Fungal Orthogroups website. OB and RNase II domains were annotated based on information from SGD (Engel et al., 2025). The NLS was previously defined by (Kurischko, Kuravi, et al., 2011) at residues 417-427, which agreed with the prediction from (Timmers et al., 1999). The NES was determined at residues 1170-1179-LSKELSDLHL, according to guidelines provided by (la Cour et al., 2004). Predictions regarding RNA and protein contacts were based on the mouse Dis3L2 structure using Phyre 2.2 (Powell et al., 2025). Rationale for other mutation constructs is found in Table 1. Domain annotations are available in Dataset 1.

Mutant proteins were first cloned onto a CEN plasmid and then integrated into each strain’s genome as follows. *SSD1* with ∼900 bp native upstream sequence was cloned with an in-frame, C-terminal GFP sequence followed by the native *SSD1* 3’UTR, onto CEN plasmid pRS-41 (derived from pRS-413 (Sikorski & Hieter, 1989)) in which the *HIS3* marker was replaced with the HYG-MX cassette (McCusker, 2017). All mutations were introduced into this wild-type construct. Single-residue mutations were generated using quick-change mutagenesis, and larger-scale deletions or combined distant point mutations were generated by PCR sewing of desired fragments, followed by Gibson cloning into pRS-41. After sequence verification, PCR fragments spanning *ssd1::GFP* and native upstream and downstream sequences were integrated into the native *ssd1::KAN-MX* locus in strain AGY1599 (haploid YPS1009 *ssd1::KAN-MX*), by co-transforming with a plasmid encoding CrispR-Cas9 and encoded guide RNA targeting the *KAN-MX* cassette, as previously described (Myers et al., 2019). Correct integration was verified by diagnostic PCR and locus sequencing. Cells were passaged overnight in YPD, and clones verified to have lost the plasmid marker were stocked and stored at -80C. The KANMX cassette was also deleted with CRISPR and verified in strains AGY1603 and 1604 to produce markerless *ssd1Δ0* knockout controls. YPS1009_Chr12 aneuploid versions of all mutants were made by crossing each mat a euploid with a mat alpha version of the YPS1009_Chr12 aneuploid (AGY1597). In all cases, Chr12 duplication was verified by diagnostic PCR as described in (Hose et al., 2020) and flow cytometry as described (Chen et al., 2012).

### Western blot analysis

Western blots characterized ssd1-GFP signal expressed in euploid and YPS1009_Chr12 aneuploids grown side-by-side and run side-by-side on SDS page gels. Westerns were exposed using anti-GFP (Abcam ab290) antibody at a 1000X dilution and anti-PGK (Abcam ab113687) at a 10,000X dilution in Licor buffer, incubated for one hour at room temperature. Westerns were developed with goat anti-mouse (Licor IRDye800CW) and goat anti-rabbit (Licor IRDye680LT) secondary antibodies, quantified in a Licor Odyssey Infrared Imager. GFP signal was quantified on the Licor system and normalized to Pgk1 signal in each lane.

### Viability with cbk1Δ

A *CBK1* knockout strain was generated by homologous recombination in a strain lacking *SSD1* (AGY1680). Viability of each mutant in the context of *cbk1Δ* was assessed by transforming AGY1680 with plasmids carrying each mutant. Mutants with wild-type levels of viable transformants were called as viable, whereas those producing no transformants were scored as not viable.

### Microscopy

We experimented with several protocols to induce Ssd1-GFP foci and found that the following produced the greatest number of visible foci in wild-type YPS1009_Chr12 cells: Strains were grown to 0.5 OD_600_ in synthetic complete media containing 2% dextrose and 2% methyl-alpha-D-mannopyranoside to minimize clumping (Sigma Aldrich) at 30°C with continuous shaking. Cells were pelleted at 5K rpm for 2 minutes, rinsed once with and then resuspended in medium without glucose, and incubated overnight at 30°C with continuous shaking. Cells were fixed in 37% formaldehyde (Sigma Aldrich), then resuspended in 0.1M KPO_4_ for imaging on poly-lysine coated sides. Images were acquired as z-stacks using an EVOS FL Auto 2 with a 100X Nikon oil immersion objective, using bright field and EVOS GFP light cube. Nuclear distribution of ssd1-GFP mutants was interrogated by visual assessment of z-stacks in NLS and NES mutants, which showed clear GFP signal in both compartments. Foci were scored manually, in part because foci of different character and size may be counted differently by automated scoring. Z-stacks were compressed and fields of ∼50 cells were assessed semi-quantitatively based on: number of cells with visible foci and average number of foci within them; amount of focus signal versus cytoplasmic signal, notes on foci size and shape. Scores are available in Dataset 3.

### RNA binding assays

Euploid mutants were grown to a 0.6 OD_600_ in YPD media, and 30 OD units were harvested by centrifugation for the RIP assay (RNA immunoprecipitation). Cells were washed twice with 1X TBS buffer and resuspended in 50 μL Ribonucleoside Vanadyl Complex (New England Biolabs) with 0.5 mL FA lysis buffer (50 mM HEPES, 140 mM NaCl, 1 mM EDTA, 1% Triton-X, 0.1% sodium deoxycholate) containing 0.125 units RNase inhibitors (Promega) and 2X protease inhibitors (Thermo). Resuspended cells were vortexed on Mini Bead Beater 16 (BioSpec Products, Model 607) with washed zirconia beads for 3 cycles of 3 minutes (with 1 minute rest between cycles) at 4°C. Lysate was treated with RQ1 DNase (Promega) for 15 minutes at room temperature and quenched with 20 mM EDTA. Lysate was centrifuged at 16g for 10 minutes at 4°C, transferred to a clean tube, then centrifuged for another 5 minutes to remove cellular debris. This was treated as the Input to the immunoprecipitation. Input was diluted 2:3 with dilution buffer (FA buffer without the detergents) and transferred to 25 μL washed GFP-Trap Magnetic Agarose beads (Proteintech Group Inc) to rotate end-over-end for 4 hours at 4°C. GFP-trap beads were washed with 230 μL (1X) and 110 μL (3X) volumes of dilution buffer then eluted in 50 μL RNase-free Laemmli sample buffer plus protease inhibitors at 70°C for 10 minutes. RNA from Input and IP was purified using RNeasy MinElute Clean Up kit (Qiagen). RNA (2 μg for INPUT and ∼500 ng for IP) was reverse transcribed to cDNA using both oligo dT and random hexamer primers with SmartScribe reverse transcriptase. (Takara). cDNA was purified using QIAquick PCR Purification kit (Qiagen) and quantified with Qubit ssDNA Assay kit (Invitrogen Q10212). Quantitative PCR (qPCR) analysis was performed using the LightCycler 480 II (Roche Diagnostics) and the LightCycler 480 SYBR Green 1 kit (Roche Diagnostics). The primer sets used for qPCR analysis are as follows: *SUN4* Forward 5’-AAC TTT GGC GCT GGT TCT TC-3’, *SUN4* Reverse 5’-TCA TCA GCG GCG ACA ATT TT-3’, *ACT1* Forward 5’-AAG GAA ATC ACC GCT TTG GC -3’, *ACT1* Reverse 5’-GGA AGG TAG TCA AAG AAG CC-3’. qPCR values for *SUN4* were normalized to *ACT1* (ΔCT) and IP values were normalized to INPUT (ΔΔCT); values are expressed as 2^^(-ΔΔCT)^.

## Supporting information

Table S1: Description of strains and mutations used in the manuscript.

Dataset 1: Snapgene file of Ssd1 domains

Dataset 2: phenotypes used for correlation analysis in Figure 5

Dataset 3: granule foci scoring

## ACKNOWLEDGEMENTS

We thank James Hose and members of the Gasch Lab for constructive discussions.

## FUNDING

This work was funded by a grant from the National Institutes of Health R01GM148975.

## SUPPLEMENTAL DATA

**Table 1:** Strains and mutant details.

**Dataset 1:** Snapgene sequence file annotating Ssd1 protein sequence domains.

**Dataset 2:** Data for phenotype correlations in Fig 4. Each column represents the phenotype and source. Growth rates represent the average of replicated growth rates in media as indicated.

**Dataset 3:** Semi-quantitative scores for ssd1-GFP foci. Mutant scores were determined manually by assessing across scores shown including: number of fields with any foci, # foci per cells in those with foci, background signal similar to ssd1- or WT control, notes on foci

## SUPPLEMENTAL FIGURES

**Figure S1.**
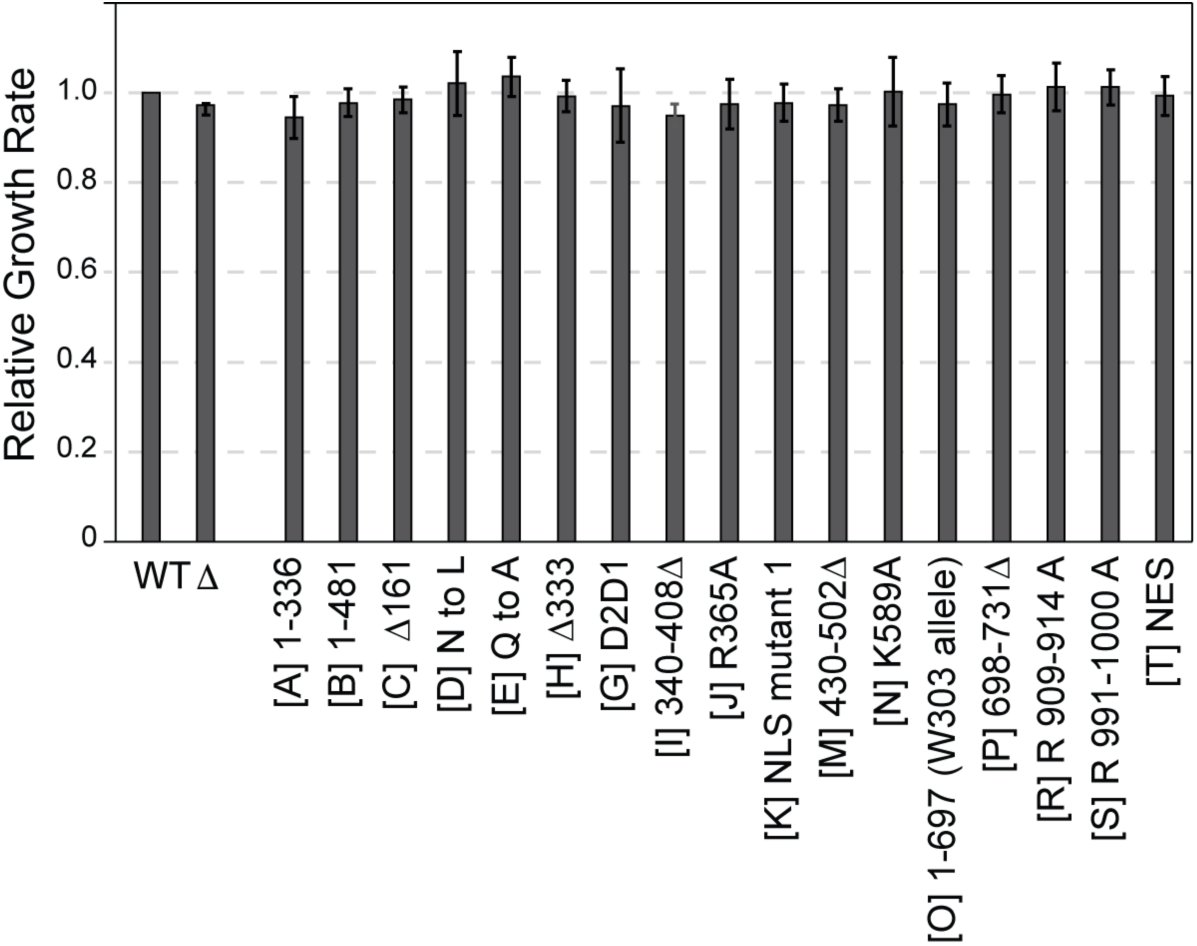
Growth rates of ssd1 mutants expressed in euploid YPS1009. Average and standard deviation (n =3) of growth rates for denoted mutants, normalized to replicate-paired wild-type. All p-values > 0.05, replicate-paired T-test.

**Figure S2.**
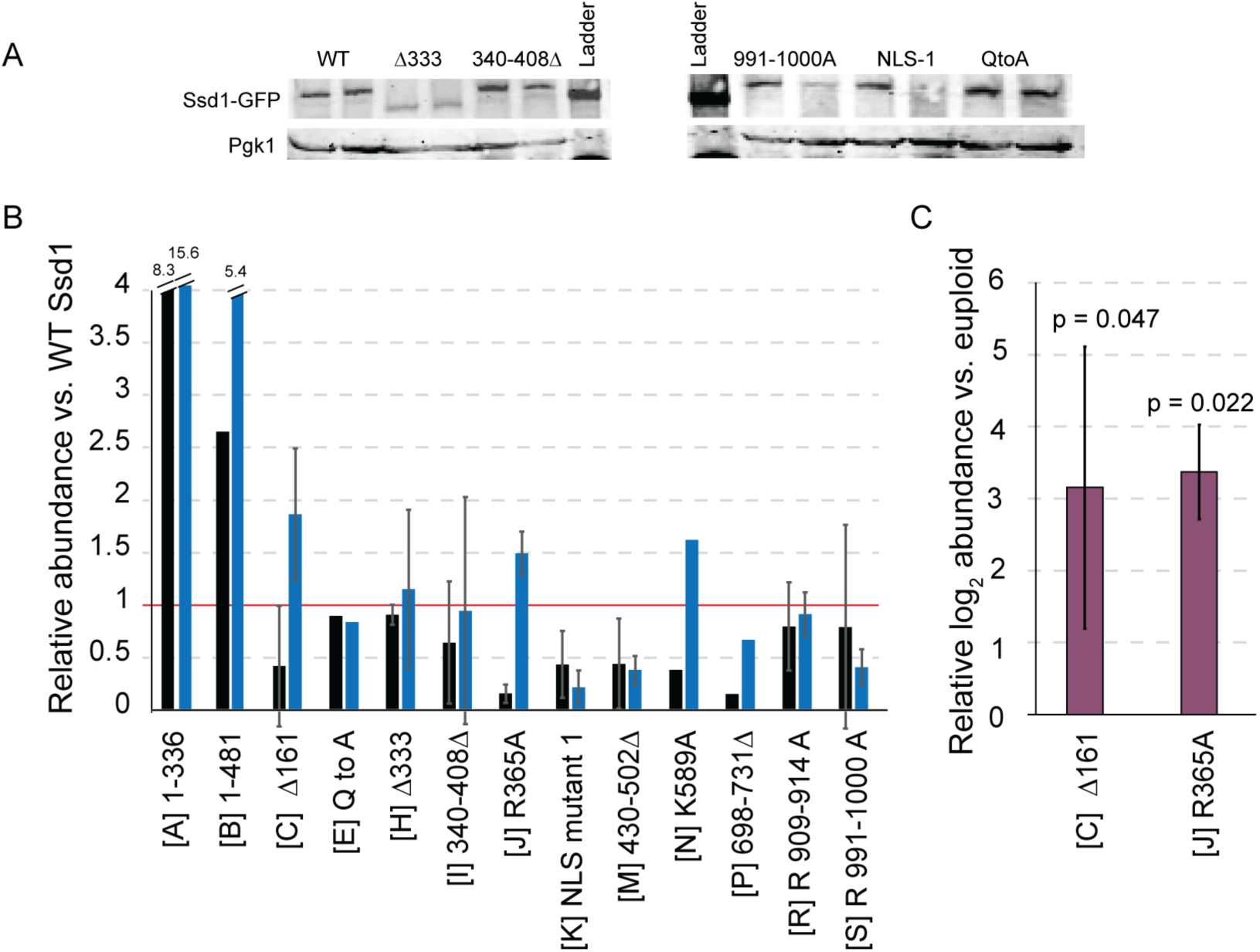
Relative abundances of specific mutants. A) Representative Westerns quantifying each Ssd1-GFP mutant in euploid (first of each annotated lane pair) and Chr12 aneuploid (second of each lane pair), run side by side and normalized to Pgk1. B) Average and standard deviation (n = 1-3) ssd1 abundance for denoted mutants expressed in euploid (black) and YPS1009_Chr12 aneuploid (blue) cells, normalized to Pgk1 loading control and plotted relative to the normalized wild-type Ssd1-GFP levels. C) Average and standard deviation (n = 3) for two mutants expressed significantly higher in YPS1009_Chr12 versus euploid cells. As described in A, plotted for aneuploid versus replicate-paired euploid. p-values (replicate-paired T-test) indicated above each plot. Several mutants were reproducibly lower in abundance than wild-type in both strains, including mutant K, M, P, and possibly S. Mutants A and B were reproducibly expressed much higher in both euploid and aneuploid cells. Mutants C and J were reproducibly lower in euploid but much higher in aneuploid cells, suggesting feedback regulation that potentially complements partial defects in these mutants (see main text).

**Figure S3.**
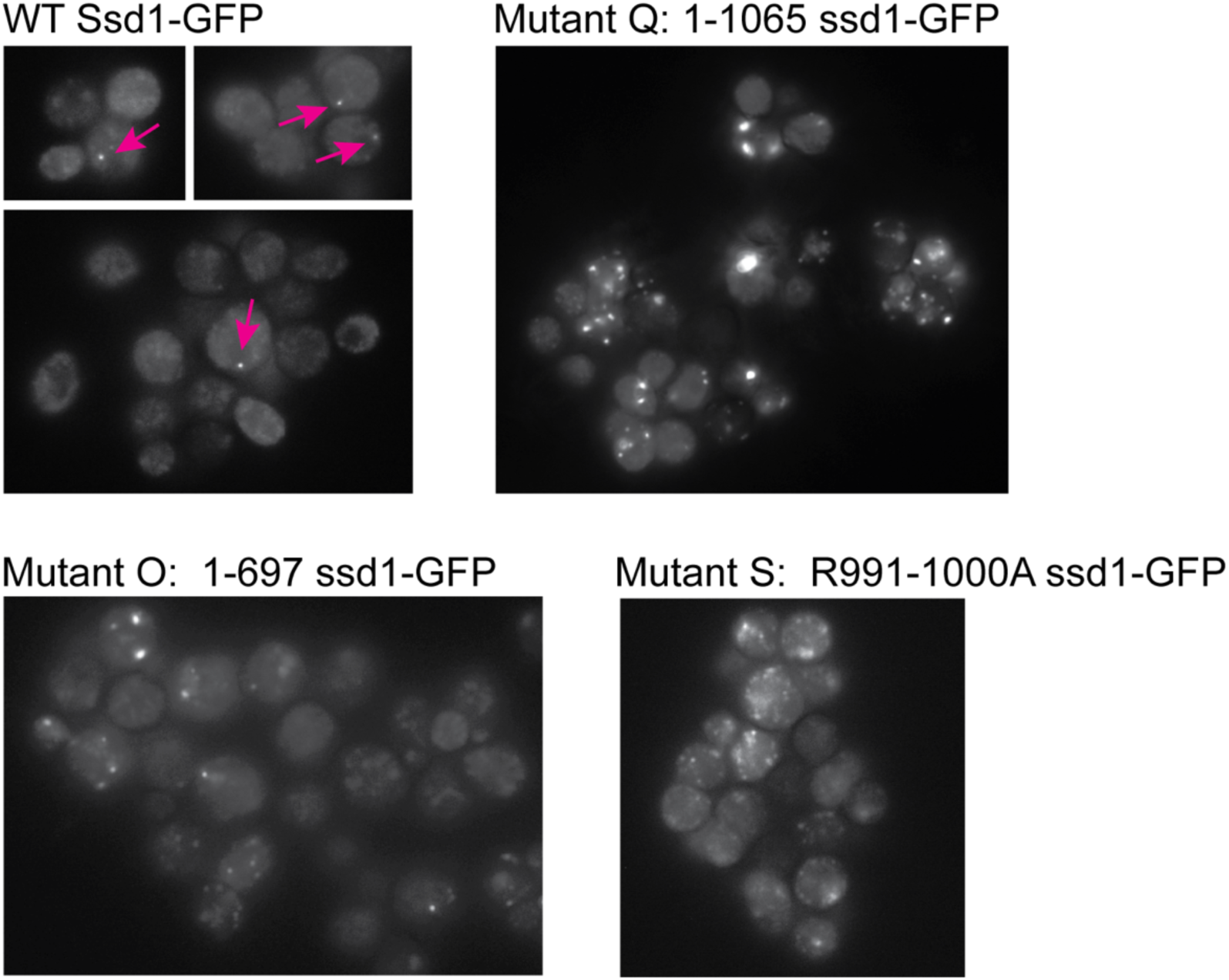
Representative Ssd1-GFP foci. Pink arrows indicate normal foci seen in euploid cells carrying Ssd1-GFP.

**Fig S4.**
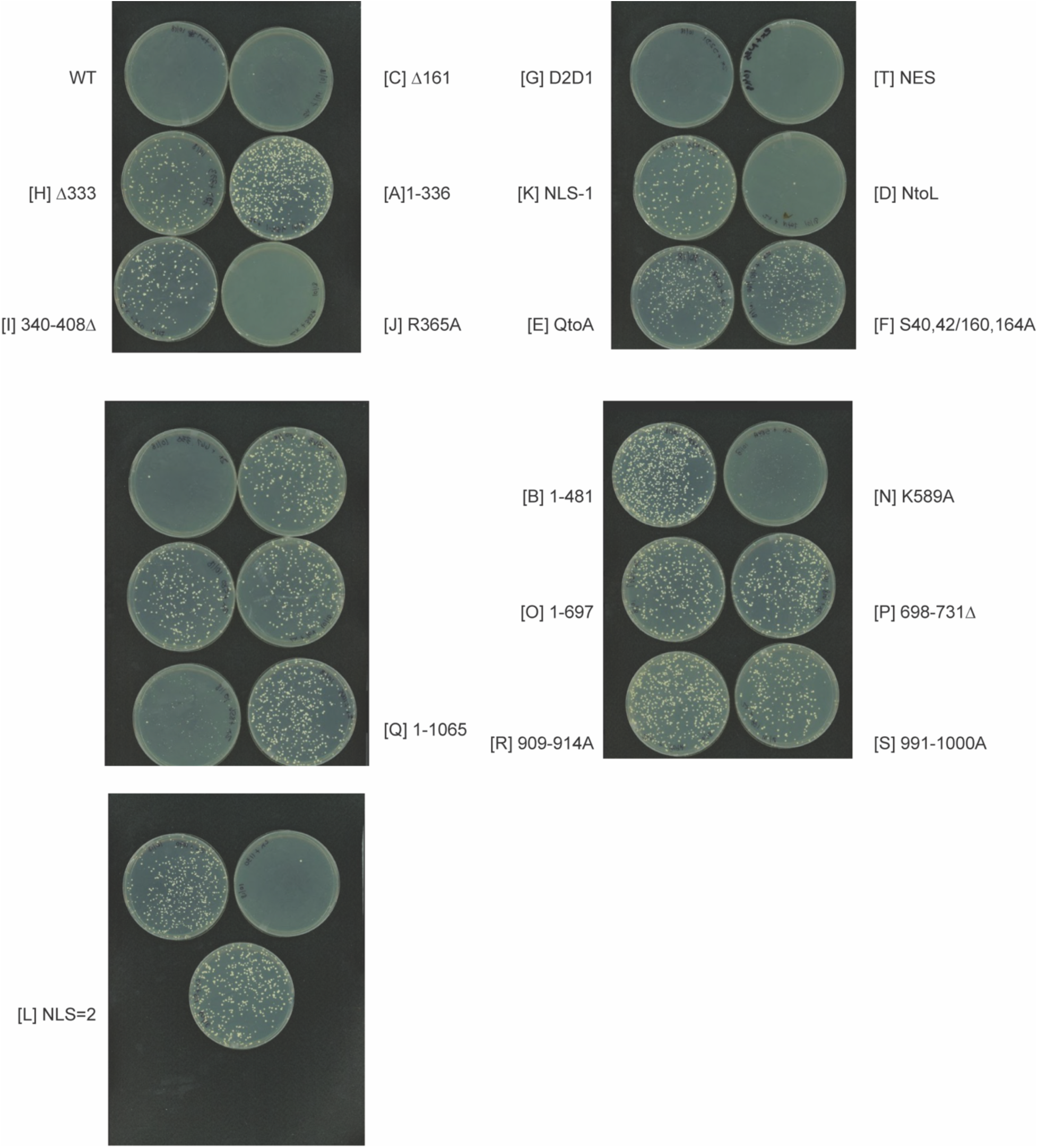
*cbk1Δ* viability assays. Representative transformation plates showing viability of denoted mutants expressed in YPS1009 *cbk1Δ* euploid cells.

## Notes

### Competing Interest Statement

The authors have declared no competing interest.

